# The magnetic field-dependent fluorescence of MagLOV2 in live bacterial cells is consistent with the radical pair mechanism

**DOI:** 10.64898/2026.02.18.706690

**Authors:** Brian L. Ross, Alessandro Lodesani, Clarice D. Aiello

**Affiliations:** Quantum Biology Institute, Los Angeles, USA

## Abstract

MagLOV2 is an engineered flavoprotein designed to have large changes in fluorescence intensity in response to weak magnetic fields. Here, we characterize the magnitude of these fluorescence changes, known as the “magnetic field effect,” as a function of the strength of an externally applied magnetic field in *E. coli* colonies expressing MagLOV2. We observe that the magnetic field effect is positive at low magnetic fields, reaches a maximum positive value near 1 mT, and then decreases, reversing sign at approximately 2 mT. Furthermore, the effect starts to plateau above approximately 70 mT, with a decreased sensitivity of fluorescence changes to magnetic fields above this range. The non-monotonic behavior, as well as the diminished responsiveness to higher magnetic fields, are consistent with the changes in fluorescence being driven by electron spin-dependent chemical processes governed by the radical pair mechanism.

## 1. Introduction

Flavoproteins have been shown *in vitro* to sense weak magnetic fields through a quantum phenomenon known as “the radical pair mechanism”. In this mechanism, light excites an electron on the flavin fluorophore, which in turn triggers an electron transfer from a nearby electron donor [1]. This transfer generates a spin-coupled radical pair in a superposition between triplet and singlet states, which an applied external magnetic field can modulate. Such an effect was first studied in depth in cryptochrome, a protein involved in regulating circadian rhythms and that is hypothesized to be involved in the magnetoreception of migratory birds [2, 3]. Sensitivity to weak magnetic fields has also been demonstrated in a handful of other flavoproteins, such as photolyase [4].

Recently, a flavoprotein known as MagLOV, along with its improved variant MagLOV2, was engineered to exhibit a large magnetic field effect (MFE), or large changes in fluorescence intensity in response to external magnetic fields [5, 6]. Derived from the LOV2 domain of a plant phototropin from *Avena sativa* (oat), MagLOV2 has been developed as a testbed for magnetogenetic tools. This protein was engineered through rounds of directed evolution, which resulted in mutations that, despite disrupting the native function of the protein, changed the chemical environment of the FMN fluorophore and enhanced magnetosensitive modulation of its fluorescent output [5]. However, the quantitative relationship of its magnetic field-dependent fluorescence response as a function of magnetic field strength has not yet been characterized. In this micropublication, we address this gap and gauge whether the observed relationship is compatible with the quantum model of the radical pair mechanism.

## 2. Methods

### 2.1. Expression of MagLOV2 in *E. coli* Colonies

The plasmids pRSETb (empty vector control) and pRSET-MagLOV2 (generously supplied to us by Maria Ingaramo from Nonfiction Biosciences) were transformed into *E. coli* BL21(DE3) cells through heat shock (or streaked from a glycerol stock), plated onto Lysogeny Broth (LB) agar plates with 100 µg/ml ampicillin or carbenicillin, and incubated overnight (Fig. 1A). A colony was picked into liquid culture, grown to mid-log phase (approximately 3 hours), and diluted approximately 10^4^-10^5^ times before plating, so that colonies were well separated. They were plated onto LB agar plates with 100 µg/ml ampicillin, 0.5 mM IPTG, and 1 µM riboflavin. Colonies were grown at 37°C overnight and then incubated at room temperature in the dark for at least another day before imaging.

**Fig. 1:**
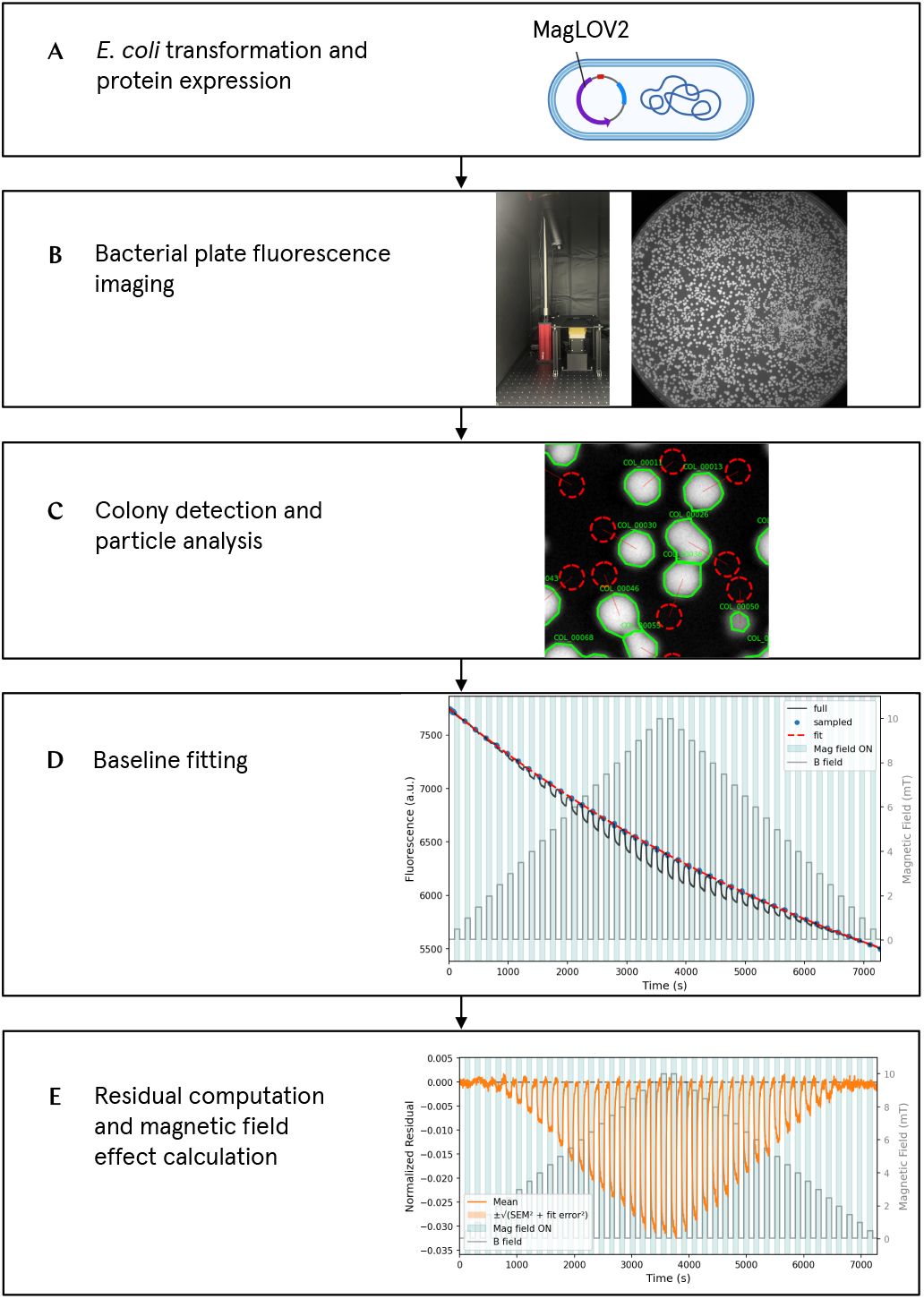
Experimental and analysis pipelines for isolating magnetic field effects in fluorescence emitted from proteins expressed in bacterial colonies. A) *E. coli* BL21(DE3) cells are transformed with an expression vector cloned with MagLOV2 or without, and plated on a bacterial plate with antibiotic for selection. B) Fluorescence images over a time-lapse are captured using the “Bacterioscope”, a bacterial plate fluorescence imaging system composed of LED illumination and detection with an sCMOS camera, with precise control of magnetic field conditions with a vector electromagnet. C) These images are automatically segmented to isolate colonies (green outline). Background regions of interest were chosen nearby to each colony for background subtraction (red rings). D) Fluorescence data from external magnetic field OFF segments are fitted with a multi-exponential model to represent the fluorescence trace expected without applied magnetic fields. E) Magnetic field effects are isolated by calculating the normalized residual of the fluorescence trace with respect to the multi-exponential fit.

### 2.2. Fluorescence Imaging of Bacterial Plates

A custom-built magneto-fluorescence imaging platform (“Bacterioscope”) was used to apply controlled magnetic field sequences during fluorescence imaging (Fig. 1B). The system combines a three-axis, water-cooled electromagnet beneath the sample stage with synchronized LED excitation at 470 nm and an externally triggered imaging with an sCMOS camera. Magnetic fields, illumination timing, and image acquisition are synchronized via hardware-timed control using a National Instruments DAQ and Python-based software. Magnetic field values at the sample plane were carefully calibrated using a magnetometer to ensure reproducible field conditions across experiments. The magnetic field used in the experiments performed in this study was arbitrarily chosen to be always out of the petri dish plane (vertical) and pointing up. The “zero applied” field condition includes the residual magnetic field when the electromagnet is off, which corresponds roughly to the Earth’s magnetic field (46 µT, 11.4° declination in Los Angeles, California) [7], with a small additional, non-uniform component (roughly *±*20% Earth’s field in amplitude) created by magnetized elements in the electromagnet. Applied magnetic field OFF and ON segments were alternated for equal periods of time (either 45 s or 90 s), such that the photophysics of the system reached a steady state by the end of the segment. For detailed information on the Bacterioscope, visit this electronic lab notebook excerpt: https://data.quantumbiology.org/bacterioscope-build-notebook.

### 2.3. Data Analysis

Fluorescence images of bacterial plates were processed using the custom-developed mfe-fit software package, which is freely available in a GitHub repository: https://github.com/Quantum-Biology-Institute/mfe_fit_analysis. Images were automatically segmented into colonies using rolling ball background subtraction, followed by thresholding with the Otsu algorithm with watershed separation (Fig. 1C). For each colony, the mean fluorescence intensity was measured and the background fluorescence from a nearby region was subtracted. The data were divided into magnetic field ON and OFF segments. To isolate the MFE, the last five data points from all OFF segments of the fluorescence trace of each colony were fitted using one to four exponential models (Fig. 1D). The model with the lowest Bayesian Information Criterion (BIC) score was selected. The fluorescence intensity was modeled using the equation:

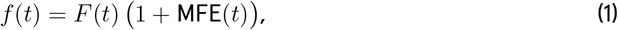

where *F* (*t*) represents the baseline fluorescence and MFE(*t*) represents the magnetic field effect. *F* (*t*) was modeled using the exponential curve fit of the OFF segments, *f*_fit_(*t*) in our package. Solving for MFE(*t*) yields:

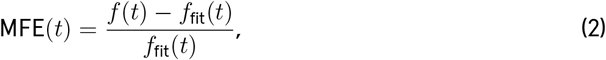

which represents the normalized fluorescence residual, or the percent deviation from the baseline. The MFE for each ON segment was computed as the normalized residual of the final five frames, corrected by subtracting the normalized residual of the final five frames from the preceding OFF segment (Fig. 1E). By taking this difference, we compensated for imperfections of the fitting (such as those observed in Fig. 3B and 3D).

MFE values were aggregated using a three-level hierarchical approach. First, all ON-OFF cycles at the same magnetic field strength within each colony fluorescence trace were averaged to obtain a per-colony mean MFE. These colony means were then averaged across all colonies within a plate, and finally an overall average was calculated across plates.

Uncertainties were propagated through each level of this hierarchy. For individual MFE measurements, fit errors were derived from covariance matrices of the nonlinear least-squares baseline fits. At the colony level, the uncertainty was calculated by taking the square root of the sum of squares of the standard error of the mean across ON-OFF cycles and the fit uncertainty. At each subsequent level of aggregation, the total uncertainty combined two components: the standard error of the mean and the propagated uncertainty from the level below.

## 3. Results and Discussion

To investigate the relationship between MagLOV2’s fluorescence and magnetic field intensities, *E. coli* BL21(DE3) bacterial plates expressing pRSET-MagLOV2 were imaged using the Bacterioscope. An external magnetic field ranging from 0.5 to 2.5 mT was applied in alternating ON and OFF segments of 45 s, the latter corresponding to geomagnetic conditions (~50 µT) (Fig. 2). Fluorescent images were captured every second. At 0.5 mT and 1.0 mT, a positive MFE, defined as an increase in fluorescence in response to the applied magnetic field, was observed. At 1.5 mT, little MFE was detected. In contrast, at 2.0 mT and 2.5 mT, a negative MFE was observed, indicating a decrease in fluorescence upon field application (Fig. 2A). This change in the sign of the magnetic field-induced fluorescence response is also evident in representative fluorescence traces from the same single representative colony, with fluorescence increasing upon field application at 1.0 mT and decreasing at 2.5 mT (Fig. 2B).

**Fig. 2:**
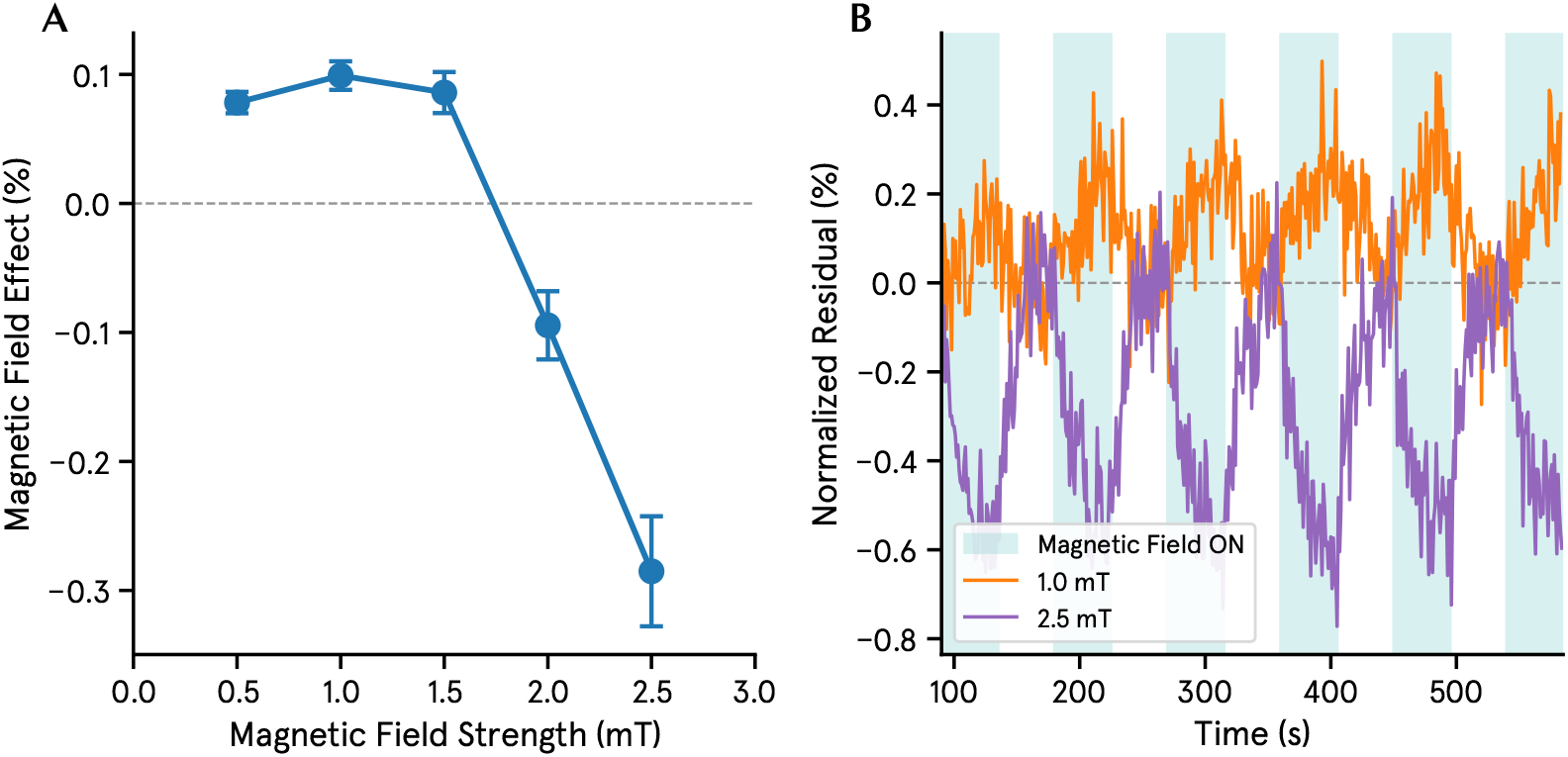
The sign of the magnetic field effect flips as magnetic field strength varies at low fields, consistent with a quantum model of the radical pair mechanism. A) Mean calculated magnetic field effects for MagLOV2-expressing *E. coli* with applied magnetic fields at 0.5 mT, 1.0 mT, 1.5 mT, 2.0 mT, and 2.5 mT (one experiment at each condition, 74 colonies each). The error is expressed as the standard error of the mean, with fit error, variation of cycles, and biological variation propagated. B) Representative fluorescence traces of the same colony for 1.0 mT and 2.5 mT. The data are from a single plate with 74 colonies. A positive magnetic field effect in fluorescence upon application of the magnetic field is observed at 1.0 mT, whereas the effect is reversed to negative at 2.5 mT. The dashed line in both panels indicates the geofield baseline.

We then performed experiments in which the applied magnetic field was ramped to obtain a more complete picture of the MFE as a function of the magnetic field strength. The experiment consisted of repeated ON-OFF cycles of an applied external magnetic field (90 s ON, 90 s OFF), with image acquisition every 2 s. During each ON phase, the magnetic field was ramped either from 0.5 to 10 mT in 0.5 mT increments or from 5 to 100 mT in 5 mT increments (Fig. 3A-D). The experiments were repeated under two conditions: with the ascending ramp preceding the descending ramp, and with the descending ramp preceding the ascending ramp.

**Fig. 3:**
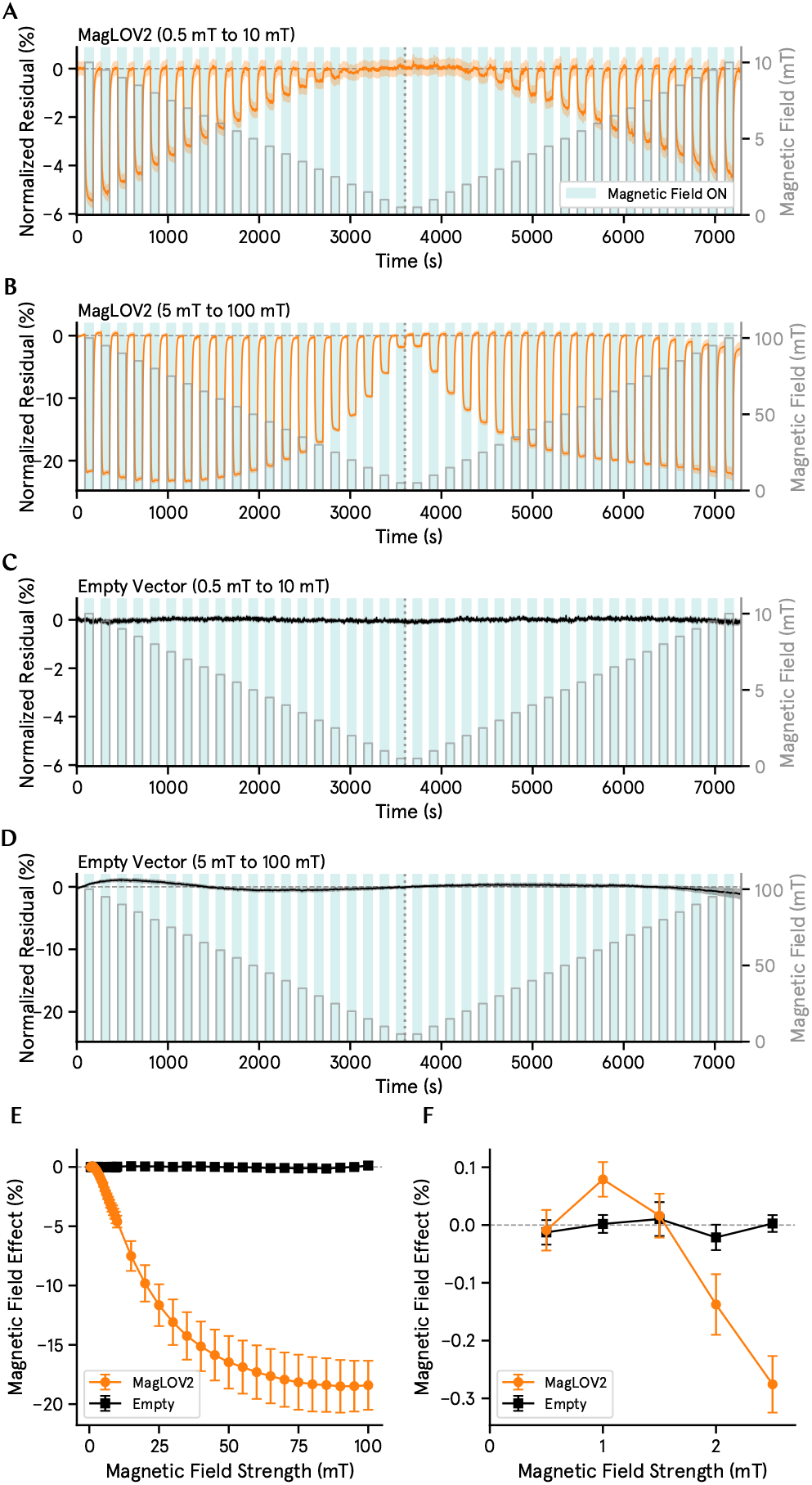
The magnetic field effect as a function of external magnetic field strength is non-monotonic at low fields and exhibits diminished sensitivity to changes in magnetic field at high fields. A-D) Representative traces of normalized residuals of bacterial colony fluorescence, isolating the magnetic field effect, for MagLOV2 (A,B) and empty vector (C,D), with ramped applied external magnetic field. Curves represent the mean across colonies for that representative experiment, with shading representing the standard error of the mean, with fit error propagated. E-F) Magnetic field effect vs. magnetic field strength relationship. Data is compiled from four plate replicates of bacterial colonies containing MagLOV2 for each magnetic field range, and three plate replicates for the empty vector. An expanded view of the low-field range (F) demonstrates a maximum positive magnetic field effect at 1.0 mT and a sign change at 2.0 mT.

Displaying the same trend as the previous set of experiments, the MFE was positive between 0.5 mT and 1 mT and increased with increasing magnetic field strength (Fig. 3E, and an expanded view 3F). Above 1.0 mT, however, the MFE decreased with increasing field strength, crossed zero at roughly 1.5 mT, and became negative at higher fields. Beyond approximately 70 mT, the MFE no longer varied strongly with magnetic field strength and instead approached a plateau, indicating a reduced sensitivity to further increases in field strength for magnetic fields up to 100 mT.

Overall, we observe a non-monotonic relationship between the MFE and the magnetic field strength at low fields, as well as a reduced sensitivity of the MFE to increasing magnetic fields at higher field strengths. Similar profiles of MFE as a function of magnetic field strengths have previously been observed in FMN [8, 9], while the plateau at higher fields has also been demonstrated in mScarlet3 [10]. Such a profile is consistent with a quantum model of magnetosensitivity based on the radical pair mechanism.

In this mechanism, the applied magnetic field modulates the singlet-triplet superposition population of a spin-correlated radical pair, with one radical belonging to the flavin cofactor and the other to an electron donor [1–3]. The modulation of the superposition, and subsequent shift in population after spin superposition coherence is lost, is driven by the fact that, in addition to the applied external magnetic field, each electron also experiences a distinct local magnetic field from nearby nuclear spins, caused by hyperfine interactions. These differences in the local magnetic field environment from nearby nuclei lead to differences in the rate of Larmor precession for the two electrons in the spin-coupled pair, which effectively makes the electrons oscillate between a triplet and a singlet state. Notably, the extent of this mixing exhibits a non-monotonic relationship with the strength of weak externally applied magnetic fields [11].

At a true zero applied field (i.e. without the magnetic field of the Earth), several spin states have approximately the same energy. When a tiny magnetic field is applied, the degeneracy breaks and opens pathways inaccessible at zero field, resulting in increased singlet-triplet mixing [12–14]. However, increasing magnetic field strengths also leads to Zeeman splitting of the three possible triplet states: T_0_, T_+_ and T_−_. The T_+_ and T_−_ states become increasingly higher and lower in energy than the T_0_ state, respectively, making them increasingly harder to access [13]. At some point, then, this effect switches the extent of singlet-triplet mixing from increasing to decreasing. Typically, at tens of mT when Zeeman splitting dominates the hyperfine interaction, the extent of the singlet-triplet interconversion plateaus.

Specifically, the observation that the MFE increases at low fields and decreases at higher fields in MagLOV2 is consistent with a radical pair born in a triplet state. For a triplet-born radical pair, increased singlet-triplet mixing leads to an increased population in the singlet state, which relaxes to the photo-excitable ground state more efficiently and subsequently leads to an increased fluorescence intensity. In contrast, for a singlet-born ground state, in which increased singlet-triplet mixing results in higher triplet yields, we would expect an inverse response, with the MFE decreasing at low fields and increasing at higher fields [15]. In addition, a triplet-born radical pair for MagLOV2 is consistent with established photophysics of LOV and phototropin domains [6, 16, 17]. Hence, the observed relationship between MagLOV2 fluorescence intensity and applied magnetic field strength mirrors the expected relationship between singlet-triplet mixing and magnetic field strength according to established theory of the radical pair mechanism.

## 4. Conclusion and Next Steps

We demonstrated that the magnetosensitive fluorescent protein MagLOV2 has a non-monotonic MFE as a function of magnetic field strength, consistent with the radical pair mechanism. However, we only investigated magnetic field strengths greater than that of the magnetic field of the Earth. Further investigations will probe the relationship at hypomagnetic conditions as well to complete the picture. Currently, we are building a second version of the Bacterioscope, equipped with a hypomagnetic chamber, which will allow us to measure the fluorescence response down to the nT level. On the molecular biology front, mutational studies of residues around the flavin fluorophore will allow us to understand how changing the chemical environment of the flavin affects the relationship between MFE and magnetic field strength. This future work will provide insight on the molecular basis for magnetosensitivity in proteins, which, in turn, will enable a rational search for potentially magnetosensitive natural proteins with such characteristics. It will also provide knowledge of how to engineer the MFE-magnetic field relationship in flavoproteins for particular applications.

## 5. Author Contributions

Following the CRediT taxonomy [18]:

1. Brian L. Ross: conceptualization; investigation; data curation; methodology; validation; formal analysis; software; writing (original draft); and visualization
2. Alessandro Lodesani: conceptualization; methodology; investigation; software; funding acquisition; and writing (review and editing)
3. Clarice D. Aiello: conceptualization; methodology; writing (review and editing); formal analysis; software; supervision; project administration; and funding acquisition

## 6. Acknowledgements

We thank Andrew York for providing a blueprint for a bacterial plate imaging system that inspired the Bacterioscope. We also thank Maria Ingaramo and Nonfiction Laboratories for providing the pRSET-MagLOV2 construct as well as valuable insights into its expression and imaging. We would like to thank Morgan Sosa for her assistance with formatting, revising, and editing the manuscript, and we thank Michael Montague, Todd Thaxton, and Bret Gaitan for critically reading the manuscript and providing helpful feedback.

The [Quantum Biology Institute](http://quantumbiology.org/) is a California non-profit 501(c)(3) focused research organization that performs basic research underpinning the quantum biology field in an open-science fashion.

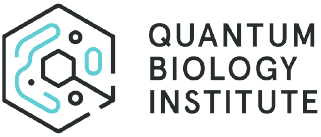

